# Rapid Assembly and Functional Differentiation of the Soil Surface Microbiome in Temperate Agricultural soil

**DOI:** 10.1101/2025.04.24.650463

**Authors:** Christopher James O’Grady, Sally Hilton, Emma Picot, Sebastien Raguideau, Christopher Quince, Christopher J. van der Gast, Hendrik Schäfer, Gary D. Bending

## Abstract

The surface of dryland soils is typically covered by biological soil crusts (BSC) dominated by cyanobacteria, lichens or moss which contribute key ecosystem functions. However, while photosynthetic biota have been observed on the surface of temperate agricultural soils, understanding of the development, biodiversity and functional significance of these communities is extremely limited. We investigated the temporal development of the soil surface community following tillage and during subsequent growth of a wheat crop. Amplicon analysis showed that establishment of the soil surface microbiome was rapid over the winter period, with distinct communities of phototrophs, bacteria and protists detected 60 days after tillage. There was rapid succession with yellow-green algae (Xanthophyceae) dominating early stages of development, followed by Cyanobacteria and Charophytes and finally moss. Temporal dynamics of fungal and protist communities was influenced by eDNA deposition from plant pathogens inhabiting the crop canopy and wild animal parasites. For all groups, assembly processes in the surface soil became increasingly dominated by dispersal limitation over time while assembly of bulk soil communities became predominantly influenced by drift. Metagenome analysis showed that the soil surface had developed a distinct functional profile relative to bulk soil after 9 months, enriched in photosynthetic processes which were largely from eukaryotic algae rather than cyanobacteria. Additionally the 3-Hydroxypropionate bicycle and reductive TCA carbon fixation pathways were enriched in surface soil. The abundance of genes involved in heterotrophic carbon, nitrogen, phosphorus and sulphur cycling processes were also enriched in surface soil. This included reads of carbohydrate active enzymes, nitrification, denitrification and sulphate reduction genes, largely attributed to the enrichment of Actinobacteria, Bacteroidetes and Proteobacteria in the soil surface. Overall, we show overlap in dominant phototrophic taxa between temperate agricultural and BSC in arid soils, but with notable differences, including abundance of Xanthphycae and Charophytes. Furthermore, the functional signatures of the soil surface communities indicate that they may play an important role in the delivery of ecosystem functions in temperate agricultural systems.

## Introduction

Complex biological assemblages, known as biocrusts or biological soil crusts (BSCs) form at the soil surface, primarily composed of phototophs, including eukaryotic algae, cyanobacteria, bryophytes, and lichens. These organisms bind and intertwine soil particles, creating a matrix that acts as a physical barrier between the soil and atmosphere (Garcia-Pichel, 2023). BSCs also support distinct heterotrophic bacterial and fungal communities (Davies et al., 2013; da Rocha et al., 2015).

BSCs are extensively distributed covering up to 12 % of the surface area of terrestrial habitats (Maier et al., 2018, Rodriguez-Caballero et al., 2018). The microbiota within BSCs play a significant role in global biogeochemical cycles, contributing to 46% of biological nitrogen fixation and 7% of net primary production in terrestrial ecosystems (Elbert et al., 2012). BSC are found on every continent and climate zone, from hyper-arid deserts to glacial forelands (Bowker et al., 2018, Belnap, 2003b, Yoshitake et al., 2010). However, most knowledge about BSC biodiversity and function comes from studies in arid and semi-arid locations.

In these ecosystems, BSCs follow a successional development pattern, with initial colonization by cyanobacteria and eukaryotic algae (Belnap, 2003b). Over time, algal dominance is replaced by mosses, lichens and liverworts. BSCs perform crucial ecosystem functions in drylands, including reducing wind and water erosion, enhancing soil stabilisation and improving water flow (Belnap and Lange, 2003). Management and engineering of BSC microbiota have been proposed as a key approach for the restoration of degraded dryland ecosystems (Coban et al., 2022).

Cropped soils used for arable cropping or permanent crops cover 11 % of the world’s surface area, and globally, agricultural soil is being eroded at unsustainable rates (Li and Fang, 2016). Agricultural soils in temperate zones may also support BSCs (Knapen et al., 2007), but unlike BSCs in drylands, understanding of their biological composition and functional significance is very limited. Knappen et al (2007) identified a succession from algal to moss dominated BSC in temperate agricultural fields, which was associated with a reduction in soil erosion. Laboratory-scale studies have shown that diverse eukaryotic and cyanobacterial phototroph communities have the potential to develop at the soil surface in temperate agricultural soils (Davies et al., 2013, Crouzet et al. 2019), but the community diversity and function of these communities *in situ* remains to be determined.

Much of the understanding of BSC microbiota is based on morphological observations. Molecular characterization of BSC community composition has largely relied on amplicon sequencing of bacterial communities, and holistic studies which integrate analysis of both eukaryotic and prokaryotic communities, as well as phototrophic and heterotrophic lifestyles are largely absent (Davies et al., 2013; Steven et al., 2012; Moreira-Grez et al., 2019).

Reliance on alpha and beta diversity analyses to characterise BSC communities has constrained the understanding of the ecological processes which shape community assembly at the soil surface during BSC development. Vellend (2010) identified four community assembly processes; natural selection, genetic drift, speciation and random dispersal. Soil surface communities may be shaped by different assembly processes compared to the bulk soil, reflecting differences in environmental selection pressures and microbial dispersal pathways. While dispersal limitation is thought to play a significant role in the assembly of BSC bacterial communities in dryland soils, the relative importance of deterministic and stochastic assembly processes is not well-defined. Factors such as moisture and nutrient availability may also influence these processes, but comprehensive studies which encompass the broader biodiversity of the BSC are lacking (Li and Hu, 2021).

Typically, microbial community assembly processes are characterised at a single point in time (Danczak et al., 2018; Chai et al., 2016). However microbial community composition shows temporal variation (Barnes et al., 2016), and it is unclear whether assembly processes remain consistent across time. This is particularly important to understand for BSCs, given the long timescales required for their formation and the dramatic changes in soil properties, morphology and associated biodiversity that occur as BSCs age.

While the importance of BSC for biogeochemical cycling in arid systems is recognized, understanding at the molecular level is poorly understood. Metagenome analysis has demonstrated that photosynthetic processes are abundant within dryland BSCs, with detection of ribulose 1,5 biphosphate carboxylase/ oxygenase (RUBISCO) and 3-hydroxypropionate cycle pathways (Steven et al., 2012; Li et al., 2020). Furthermore, various nitrogen transformation pathways have been detected in the metagenomes of dryland BSCs, including assimilatory and dissimilatory nitrate reduction, denitrification and nitrogen fixation. The relative importance of these pathways appears to vary across different types of dryland BSCs (Wang et al., 2021; Li et al, 2020). Broader functional analysis of biogeochemical cycling pathways in non-dryland BSCs remains to be investigated. Importantly, understanding the extent to which biogeochemical functions are altered at the soil surface relative to bulk soil are unclear even in dryland soils.

Here, we characterised temporal succession of biodiversity during development of soil surface communities in a temperate agricultural soil and examined how ecological processes shaping community assembly changed over time. Importantly, a holistic analysis of biodiversity was conducted by characterising bacterial, fungal and protist communities, while analysis of chloroplast 23S rRNA enabled detailed analysis of both eukaryotic and bacterial phototrophs. This was supplemented by functional comparison of soil surface and bulk soil communities using metagenome analysis.

## Methods

### Study location

Soil samples were obtained between November 2016 and August 2017 from Bradley’s field at Warwick Crop Centre, Wellesbourne, United Kingdom (52.213833°N, −1.605683°E), prior to and during growth of a winter wheat (*Triticum aestivum*) crop. The field had previously been cropped with oilseed rape (*Brassica napus*), which had been harvested in July 2016. The field was subject to 6-inch conservational tillage two weeks prior to the initial sampling date, and the wheat crop was drilled the following week. Herbicides and mineral NH_4_NO_3_ fertiliser were applied to the crop according to local best conventional practices. Rainfall, temperature and humidity data (Supplementary Fig 1 a-c) were collected from the Wellesbourne weather station, located at Warwick Crop Centre.

### Sampling methods and soil analysis

Sampling occurred at approximately 30-day intervals for ten months with the final sampling point immediately before harvest of the winter wheat crop. At each time point sampling was conducted at five positions along an east-west transect, separated by 10 m. At each position, five soil cores were collected in a W pattern with cores spaced 4 m apart. Soil cores were collected by inserting a 5 cm width x 10 cm depth stainless steel cylinder into the soil. Soil was dug from around the cylinder to retrieve it. The initial transect was located 30 metres away from the edge of the field. At each sampling time the transect was moved 10 m northwards, maintaining distance to the field edge. Cores were stored at 4°C until processing the following day.

Soil was pushed from the base of the core, and the top 2 mm of soil was removed and defined as the soil surface. This was based on preliminary research which found negligible chlorophyll below this depth in soil cores collected from the field between April-July 2016. Soil from 2 mm to 2 cm depth was removed and termed bulk soil.

BSC and bulk soil from the five samples collected at each sampling position were pooled for subsequent analysis. Bulk density was measured for each sample (Supplementary Fig 1d). Samples were sieved < 2 mm before analysis of soil moisture and pH (Supplementary Fig 1 e, f), chlorophyll α content (Ritchie et al., 2006), and microbial biomass C and N via chloroform fumigation-extraction, followed by organic C and total N analysis on a TOC-L Shimadzu analyser (Joergensen, 1996).

### DNA amplicon sequencing and bioinformatic analysis

DNA was extracted from 0.5 g of soil surface and bulk soil using the MPBio FastDNA^TM^ SPIN Kit for Soil, according to the manufacturer’s instructions. DNA was used for PCR to amplify the 16S V3 and V4 rRNA region region of bacteria (341F and 806R primers, Caporaso et al., 2011), the 18S rRNA V1-V3 region of protists (Euk-A and Euk-570, Countway et al., 2005), the ITS2 region of fungi (fITS7 and fITS4, Ihrmark et al., 2012) and the 23S rRNA region of phototrophs (rVF1 and rVR1, Sherwood and Presting, 2007). Primers were modified with Illumina Nextera Index Kit v2 adapters. PCRs were conducted using a Multigene Optimax thermal cycler (Labnet, USA), in a 25 μl volume with 15 ng DNA, Q5® Hot start High Fidelity 2x Master Mix (New England Biolabs) and 0.5 μM of each primer. Thermocycling conditions for 16S rRNA, 18S rRNA and ITS2 are described in Hilton et al. (2021) while 23S rRNA are described in Davies et al. (2013). PCR products were purified using AMPure XP beads (Beckman Coulter, Germany) according to the manufacturer’s instructions and the DNA concentrations were measured using a Qubit 2.0 Fluorometer (Life Technologies, USA) and diluted to 4 nM. The libraries were pooled and sequenced using the Illumina MiSeq Reagent Kit v300 for 2 x 300 bp sequencing.

Detailed description of sequence processing is provided in Hilton et al. (2021). Briefly, following sequencing, low-quality bases were removed using Trimmomatic v0.35 (Bolger et al., 2014). Paired-end reads were assembled by aligning the forward and reverse reads, before primers were trimmed, and quality filtering conducted using USEARCH (Edgar, 2010). Unique sequences were filtered by abundance, and singletons removed. A minimum identity threshold of 97% was used to cluster similar reads into operational taxonomic units (OTUs). QIIME 1.8 (Caporaso et al., 2010) was used to assign taxonomy of 16S rRNA to the Greengenes reference database (McDonald et al., 2012), ITS sequences to the UNITE reference database (Kõljalg et al., 2013) and 18S rRNA and 23S rRNA genes to the SILVA reference databases (Quast et al., 2013). For 16S rRNA OTUs assigned to mitochondria or chloroplasts were removed. Filtering was performed on 18S rRNA OTU tables to remove any Metazoa, Archaeplastida and fungal sequences, to leave predominantly single celled eukaryotes, hereafter termed protists. For 23S rRNA any OTUs assigned to non-phototrophic taxa were removed, leaving bacterial and eukaryotic phototrophs. Raw sequence and OTU datasets reported in this study are available in the NCBI Sequence Read Archive under BioProject ID PRJNA718625.

### Microbial diversity analyses

All OTU tables were rarefied using a single random subsampling algorithm to the lowest number of reads across all samples for each amplicon, which was 8700, 10600, 1360, 1050 reads for 16S rRNA, ITS, 18S rRNA and 23S rRNA, respectively. Rarefaction curves demonstrated effective capture of the diversity of all amplicons at this sequencing depth (Supplementary Fig 2). This resulted in 7317 (16S), 1955 (ITS), 1560 (18S) and 407 (ITS) OTUs after rarefaction.

Alpha diversity (Fisher’s alpha) and non-metric multidimensional scaling (NMDS) were calculated using the vegan package in RStudio (McMurdie and Holmes, 2013; RStudio Team, 2015) and plotted with ggplot2 (Wickham, 2016). Significant differences in Fisher’s alpha were investigated using the Kruskal Wallis test with Dunn’s test for post hoc testing, with P values corrected for multiple comparisons using the Benjamini-Hochberg method. Analysis of Similarity (ANOSIM) was used to determine statistical dissimilarity between samples at the community level (Oksanen et al., 2018). Similarity percentages (SIMPER) analysis was performed to determine which phyla, class or OTUs contributed to the differences in community structure of bacteria, fungi, protists and phototrophs in the surface soil across time.

### Community assembly processes

Microbial community assembly processes were determined according to the Vellend (2010) framework following a modified approach proposed by Stegen et al. (2013, 2015). Sequences were aligned and filtered using QIIME 1.8 which were used to create a maximum-likelihood phylogenetic tree using the muscle algorithm. The sampling time where each OTU had maximal relative abundance (referred to as the ‘temporal niche’) was determined and a Euclidean distance matrix was produced to determine temporal niche differences between each OTU, which were normalized between 0 - 1. Pairwise phylogenetic distance between OTUs was determined and were binned in intervals of 0.1. Mantel correlograms were produced using the vegan package of R between phylogenetic and Euclidean temporal distance matrices with 999 permutations (Oksanen et al., 2018).

Phylogenetic community turnover was determined by calculating β-mean nearest taxon index (β-MNTD). OTUs were randomly shuffled across the tips of the phylogenetic tree, and β-MNTD was recalculated 999 times to create a null distribution of β-MNTD and β-nearest taxon index (β-NTI). Values of |β-NTI| > 2 indicate that observed species turnover (between a pair of communities) was due to environmental selection processes, β-NTI > 2 indicates variable selection (environmental selection between two samples is significantly different), β-NTI < −2 indicates homogenous selection (Stegen et al., 2012).

For instances where no selection processes dominated (|β-NTI| < 2), stochastic processes were determined using the Raup-Crick algorithm using pairwise Bray-Curtis matrices (RC_Bray_). RC_Bray_ values > 0.95 indicate homogenising dispersal (individuals freely moving between populations), RC_Bray_ < −0.95 infer dispersal limitation (a barrier preventing movement of individuals across populations), and RC_Bray_ < 0.95 indicate drift (random changes of species abundance due to stochastic birth and death events). Pairwise β-NTI comparisons were made across all possible spatial and temporal comparisons within each compartment (within soil surface, and within bulk soil).

### Metagenomic sequencing and bioinformatic analysis

DNA isolated from soil surface and bulk soil samples collected after 270 d in August 2017 were selected for metagenome analysis. These samples were selected as they contained the most well-developed soil surface communities based on morphology, chlorophyll and biomass-C content (Fig 1a-c). 150 bp paired-end shotgun sequencing was performed on an Illumina HiSeq 4000, with a target depth of 50 Gb per sample. Raw sequences were trimmed using Trimmomatic (version 0.39) to remove 3’- and 5’-adapter sequences, low-quality base pairs, low quality reads and to merge forward and reverse reads (Bolger et al., 2014). Quality-controlled sequences from all samples (5 bulk soil and 5 soil surface) were co-assembled using MEGAHIT (version 2.0), a *de novo* de Bruijn graph based metagenomic assembler, using a *k-*mer range between 27-127, with a step interval of 10 (Li et al., 2015). Contigs below 1000 base pairs in length were removed from any further analysis. Co-assembled contigs were quality assessed using MetaQUAST using default parameters (Mikheenko et al., 2015). Raw and assembled metagenome sequence files have been deposited at the NCBI under the accession number SRR31479107.

**Figure 1.**
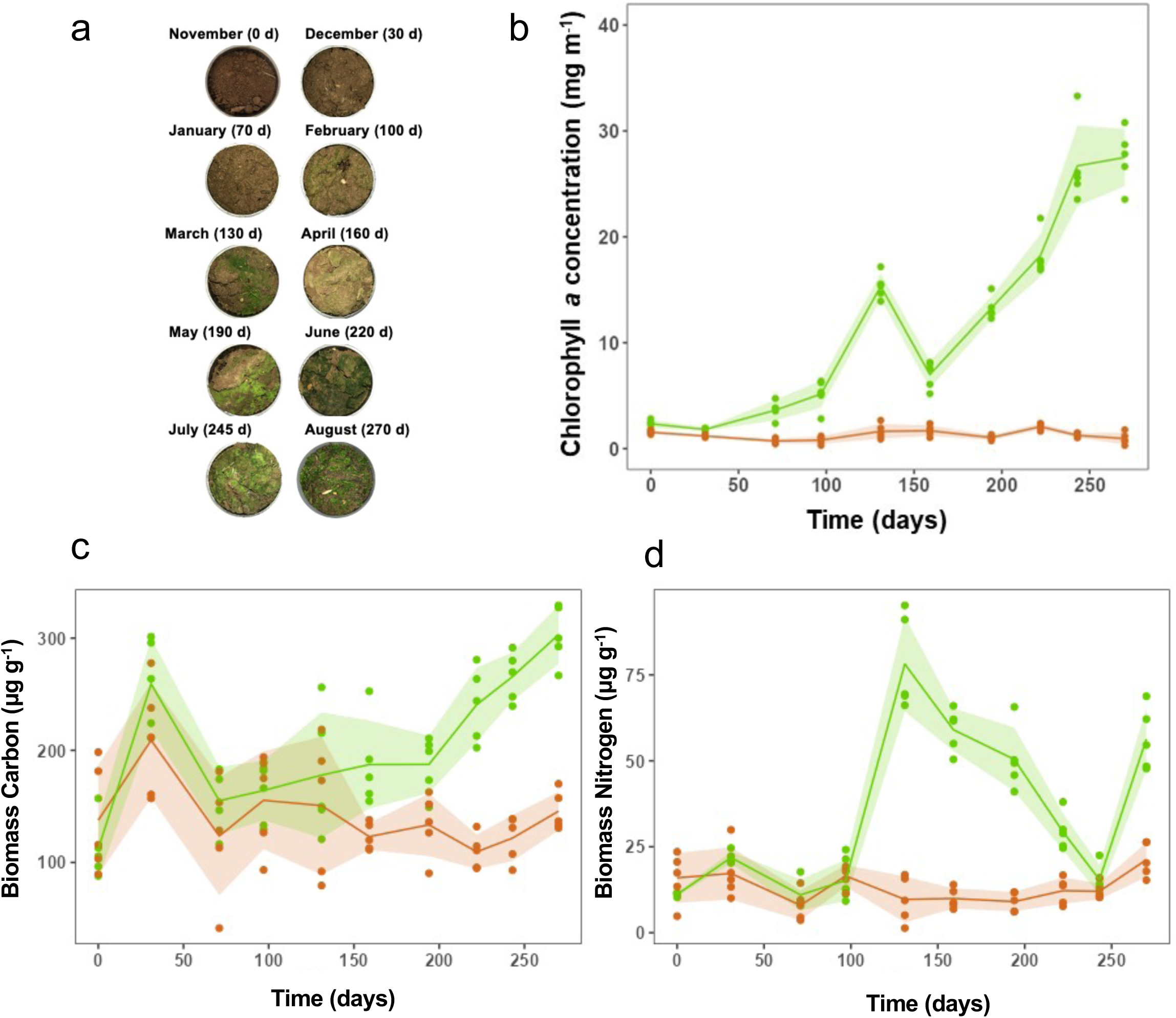
Growth of microbial communities at the soil surface over the sampling period. a. Visual appearance of the soil surface across the sampling period. Photographs of the surface of a randomly selected core taken on the day of sampling. b. Chlorophyll *a* concentration of surface and bulk soil samples. c. Biomass carbon content of surface and bulk soil samples d. Biomass nitrogen content of surface and bulk soil samples For b-d, green indicates soil surface and light brown the bulk soil. Error bars (shaded) indicate +/- standard deviation.

Open reading frames (ORFs) of co-assembled contigs were predicted using Prodigal (version 2.6.3) using the ‘meta’ pre-set (Hyatt et al., 2010). Predicted ORFs were translated into amino acid sequences and functional annotations were obtained by performing a DIAMOND search (version 0.9.9.110) which queried the Kyoto Encyclopaedia of Genes and Genomes (KEGG), Clusters of Orthologous Groups (COG), The Protein Families (Pfam) and Carbohydrate Active Enzyme (CAZy) databases using an expected value (e-value) threshold of 1e^-15^, a minimum amino acid length of 50 and a minimum identity of 90% (Galperin et al., 2014, Finn et al., 2014, Buchfink et al., 2014, Kanehisa et al., 2015, Lombard et al., 2013). Taxonomic annotations of assembled contigs were obtained using ‘CAT and BAT’, which queried contigs against The National Centre for Biotechnology Information non-redundant protein sequences database using an e-value threshold of 1e^-15^ (von Meijenfeldt et al., 2019). Quality controlled reads from each sample were mapped to annotated contigs and annotated ORFs obtained after co-assembly using Bowtie2 and subsequently filtered and sorted using SAMtools (Langmead and Salzberg, 2012, Li et al., 2009). Mapping information from each sample was collated across all samples to obtain coverage information (read hits per contig/ORF) using the ‘coverage’ function within the BEDTools suite (Quinlan and Hall, 2010). Raw coverage information was normalized using DESeq2 to adjust for sequencing depth and library size differences, incorporating soil compartment as a factor in the model (Love et al., 2014).

Tests for a normal distribution were performed using a Shapiro-Wilk test, and statistical comparisons between treatments were performed using a Wilcoxon signed-rank sum test for pair-wise comparisons. Stacked bar plots were generated using the Phyloseq package of R and were plotted using the ggplot2 R package.

## Results

### Development of a phototroph layer at the soil surface

The development of a green layer across the soil surface was visible by January (70 days) and by August (270 days) the soil surface appeared to be dominated by moss (Fig 1 a). Two-way ANOVA revealed that chlorophyll *a* concentration was significantly different between compartments (p < 0.0001), and displayed a strong, positive correlation with time in the soil surface (R^2^ = 0.8042, p < 0.0001) but not the bulk soil. There was significantly (p < 0.05) higher chlorophyll content in surface relative to bulk soil from January (70 d) onwards (Fig 1 b).

Biomass C and N significantly varied between compartments (p < 0.0001, Fig 1 c, d). with significantly higher biomass C (p < 0.05) and N (p < 0.001) in the surface compared to bulk soil from April (160 days) and March (130 days) onwards respectively. In surface soil there was a strong, positive correlation with time for biomass C (R^2^ = 0.3395, p < 0.0001) and N (R^2^ = 0.1464, p = 0.0032). No such relationships were found in bulk soil.

### Diversity and composition of soil surface and bulk soil microbial communities

When data was combined across time points, Wilcoxon signed-rank sum tests identified significant differences in Fisher’s alpha diversity between surface and bulk soil compartments for bacterial, protist and phototroph (p < 0.0001) communities (Fig 2 a i), iii), iv)), which was 1.18 (bacteria), 1.37 (protists) and 1.46 (phototrophs) times higher on average in bulk soil in comparison to the soil surface. However, there was no significant difference in Fisher’s alpha diversity of fungal communities between surface and bulk soil (Fig 2 a ii)).

**Figure 2.**
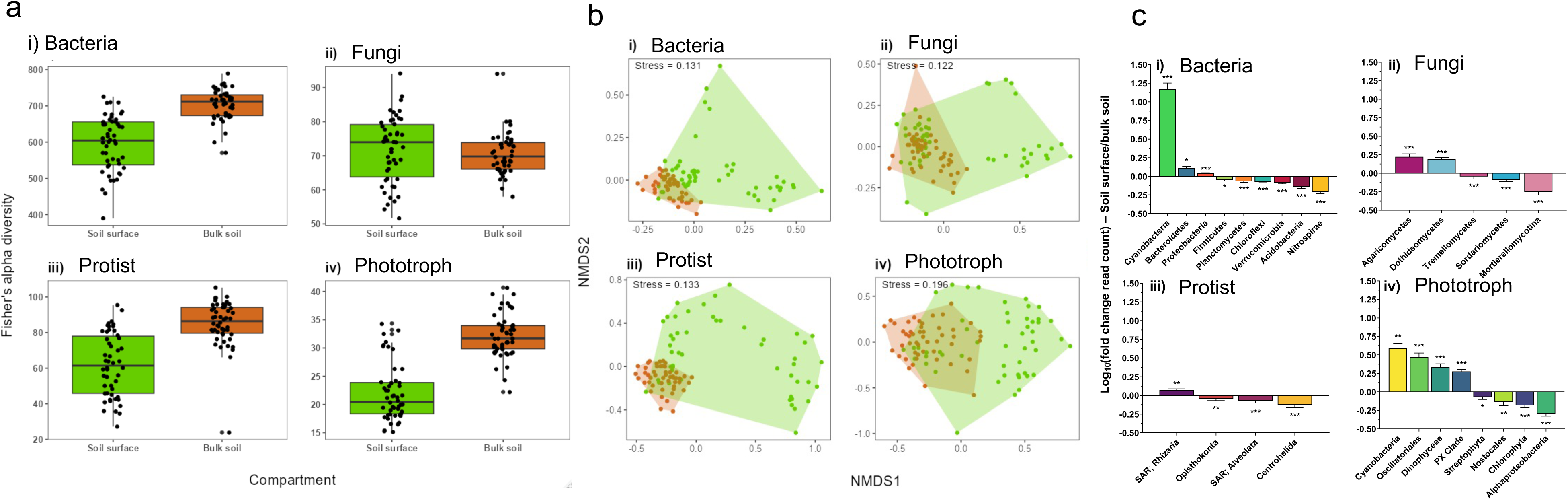
Biodiversity of bacterial, fungal, protist and phototroph communities in soil surface and bulk soil. a. Fisher’s Alpha diversity of i) Bacterial, ii) Fungal, iii) Protist, and iv) Phototroph communities in surface and bulk soil. Data is combined across all time points. b. Non-Metric Multidimensional Scaling analysis of Bray-Curtis similarity of soil surface and bulk soil biota. Data is combined across all time points. i) Bacterial, ii) Fungal, iii) Protist, and iv) Phototroph communities. c. Differential abundance of i) Bacterial, ii) Fungal, iii) Protist, and iv) Phototroph communities in soil surface and and bulk soil over the sampling period. Bacterial (16S rRNA) communities are shown at the phylum level, protist (18S rRNA) communities at the ‘ta2’ taxonomic level (phylum equivalent) and fungal (ITS) and phototroph (23S rRNA) communities at the class level. Relative abundance was transformed to fold change (relative abundance of given taxa in surface/bulk soil) and subsequently log_10_ transformed. Taxa showing significantly different abundance between surface and bulk soil using a Wilcoxon signed-rank test are indicated (*, p < 0.05; **, < 0.01; ***, p < 0.001).

When samples from all time points were combined, NMDS and ANOSIM showed there were significant differences (p < 0.001) in community composition between surface and bulk soil samples for bacteria (*R* = 0.317), protist (*R* = 0.358) and phototroph (*R* = 0.438) communities (Fig 2 b i), iii), iv)). ANOSIM revealed no significant difference between soil surface and bulk soil fungal communities, but the test statistic was close to the significance threshold of 0.2 (*R* = 0.1967).

Across all time points, Actinobacteria, Proteobacteria and Acidobacteria were the most abundant bacterial phyla in both surface and bulk soil compartments (Supplementary Fig 3 a), contributing 29.9%, 29.5 and 7.2% relative abundance in the soil surface, and 28.8%, 26.5% and 10.1% relative abundance respectively in the bulk soil. Nine of the twelve most abundant bacterial phyla showed significant differences in relative abundance between soil surface and bulk soil samples (Fig 2 c i). Soil surface samples had significant enrichments of Cyanobacteria (L_10_FC (logarithmic 10-fold abundance change in surface relative to bulk soil) = 1.168, p < 0.001), Bacteroidetes (L_10_FC = 0.111, p = 0.034), and Proteobacteria (L_10_FC = 0.041, p < 0.001). Conversely, Nitrospirae (L_10_FC = −0.207, p < 0.001), Acidobacteria (L_10_FC = −0.139, p < 0.001), Verrucomicrobia (L_10_FC = −0.087, p < 0.001), Chloroflexi (L_10_FC = −0.072, p < 0.001), Planctomycetes (L_10_FC = −0.066, p < 0.001), and Firmicutes (L_10_FC = −0.048, p = 0.037) were significantly enriched in the bulk soil.

In fungal communities, Soradiomycetes, Dothideomycetes and Eurotiomycetes were the most abundant classes in both compartments contributing to 28.5%, 21.5% and 11.47% relative abundance in the soil surface, and 35.3%, 13.8% and 11.00% relative abundance respectively in the bulk soil (Supplementary Fig 3 b). Five out of nine abundant classes displayed significant compartmental differences between soil surface and bulk soil (Fig 2 c ii)). Agaricomycetes (L_10_FC = 0.222, p < 0.001) and Dothideomycetes (L_10_FC = 0.189, p < 0.001) were significantly enriched in the soil surface, whereas Mortierellomycotina (L_10_FC = 0.255, p < 0.001), Sordariomycetes (L_10_FC = −0.093, p < 0.001) and Tremellomycetes (L_10_FC = −0.043, p < 0 .001) were enriched in the bulk soil.

In protist communities, Rhizaria (SAR), Amoebozoa and Stramenopiles (SAR) were the most abundant taxa at the phylum-equivalent level (ta2), contributing to 42.5%, 17.8% and 11.1% relative abundance in the soil surface, and 36.4%, 15.0% and 15.6% relative abundance respectively in the bulk soil (Supplementary Fig 3 c). Four out of seven phylum-equivalent taxa displayed significant compartmental differences (Fig 2 c iii)). Rhizaria (L_10_FC=-0.072, p = 0.00319) was the only taxa significantly enriched in the soil surface, whereas Centrohelida (L_10_FC = −0.123, p < 0.001, Alveolata (L_10_FC= −0.069, p < 0.001) and Opisthokonta (L_10_FC = −0.047, p = 0.0073) were significantly enriched in bulk soil.

In phototroph communities, Stramenopiles and Viridiplantae were the most abundant taxa at the phylum-equivalent level (ta2), contributing to 38.8% and 36.4% relative abundance in the soil surface, and 21.1% and 51.2% relative abundance respectively in the bulk soil (Supplementary Fig 3 d). Eight out of nine classes showed significant differences in relative abundance between compartments (Fig 2 c iv). Cyanobacteria (no class assigned, L_10_FC = 0.591, *p* = 0.0035), Oscillatoriales (Cyanobacteria, L_10_FC = 0.469, *p* < 0.001), Dinophyceae (L_10_FC = 0.336, *p* < 0.001) and the Stramenopiles PX clade (L_10_FC = 0.275, *p* < 0.001) were significantly enriched in the soil surface, whereas Alpha Proteobacteria (L_10_FC = −0.296, *p* < 0.001), Chlorophyta (L_10_FC = −0.182, *p* < 0.001), Nostocales (L_10_FC = −0.138, *p* = 0.0267) and Streptophyta (L_10_FC = −0.070, *p* = 0.0191) were significantly enriched in the bulk soil.

### Temporal dynamics of soil surface communities

In soil surface communities, regression analysis revealed that alpha diversity had a strong, significant (p < 0.0001) decline across time for all amplicons (*R^2^* = 0.3301, 0.4564, 0.5679 and 0.5475 for bacteria, fungi, protists and phototrophs, respectively (Supplementary Fig 4 a-d)). However, in bulk soil only fungal community diversity had a significant negative correlation with time although the goodness of fit of this correlation was poor (R^2^=0.121, p<0.013). Pairwise Bray-Curtis comparisons of each sample were made with samples obtained from the initial d 0 November timepoint for each amplicon. Regression analysis revealed strong significant (p <0.0001) negative correlations between Bray-Curtis similarity and time for each amplicon in both the soil surface and bulk soil. However, analysis of gradients revealed that Bray-Curtis similarity decreased at a faster rate over time in soil surface communities relative to bulk soil communities for all amplicons (p < 0.0001 in all cases).

SIMPER analysis identified key phyla (Fig 3, Supplementary Table 1) contributing to community dissimilarity over time. At early stages of soil surface community development between November (30 days) and March (130 days), Xanthophyceae (yellow-green algae, Fig 3 d) and proteobacteria (Fig 3 a) showed marked increases in relative abundance, but subsequently declined as Klebsormidiophyceae (charophyte algae, Fig 3 d), Basidiobolomycetes (Fig 3 b), Dothidiomycetes (Fig 3 b), Rhizaria (Fig 3 c) and *Microcoleus* (Fig 3 d) increased in relative abundance between January (70 days) and April-July (160-245 days). Relative abundance of these groups in turn declined over time, while relative abundance of embryophytes (which includes bryophytes, Fig 3 d), Leotiomycetes (Fig 3 b), Eurotiomycetes (Fig 3 b), Amoebozoa (Fig 3 c) and proteobacteria (Fig 3 a) increased, peaking in August (270 days).

**Figure 3.**
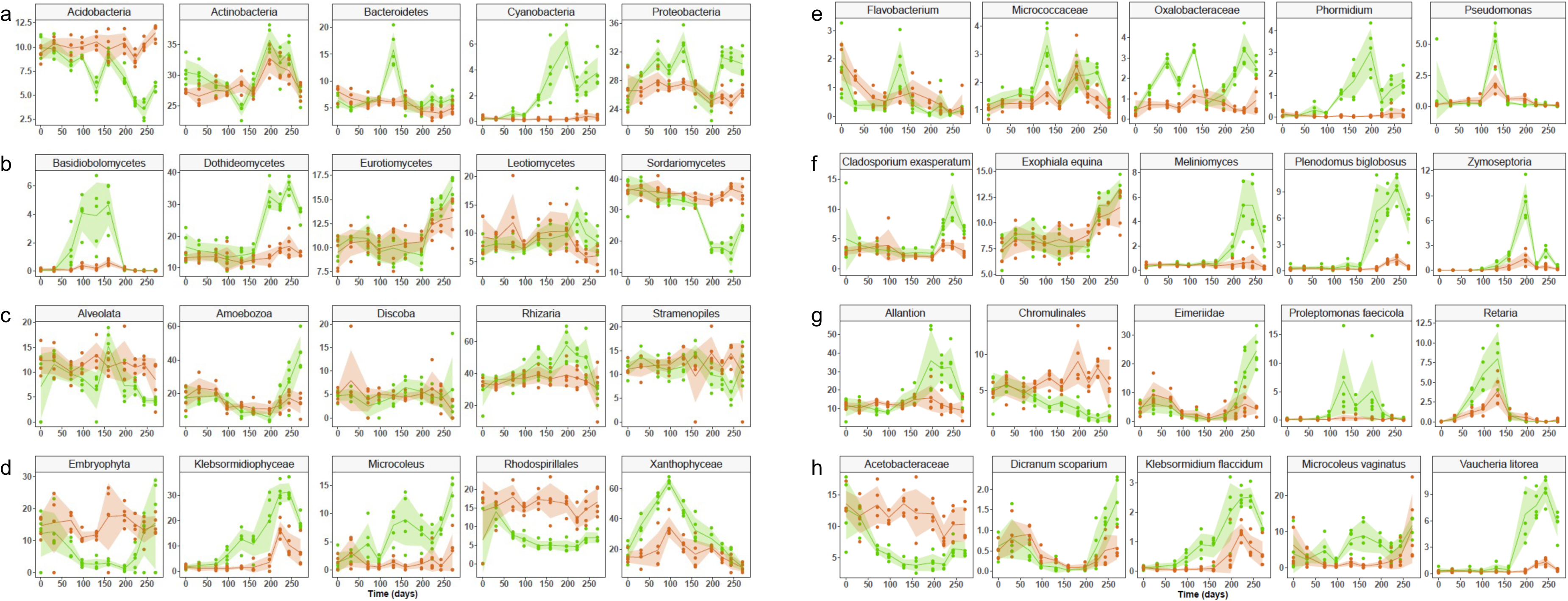
Taxonomic changes in soil surface and bulk soil bacterial, fungal, protist and phototroph communities. Key taxa contributing to community dissimilarity between surface (green) and bulk soil (light brown), as determined by Similarity Percentage Analysis (Supplementary Table 1). The % relative abundance over time of the top five taxa are shown in each case. a. Bacterial phyla, b. Protist ‘ta2’ taxonomic level, c. Fungal class, d. Phototroph class, e. Bacterial OTU, f. Fungal OTU, g. Protist OTU, h. Phototroph OTU. Shaded regions indicate +/- standard deviation.

Similarly at the OTU level, SIMPER analysis (Supplementary Table 1) identified key taxa associated with changes in community composition over time. *Phormidium* sp. (Cyanobacteria, Fig 3 e), *Pseudomonas* sp. (Proteobacteria, Fig 3 e), *Proleptomonas faecicola* (Euglenozoa, Fig 3 g), Retaria (Rhizaria, Fig 3 g) and Zymoseptoria (Dothidiomycete, Fig 3 f) showed distinct peaks in relative abundance between March (130 days) and May (196 days), while relative abundance of *Plenodomus biglobosus (*Dothidiomycete, Fig 3 f*), Cladosporium exasperatum* (Dothidiomycete, Fig 3 f), *Meliniomyces* sp. (Leotiomycete, Fig 3 f), Eiermeriidae (Alveolata Fig 3 g), *Vaucheria litorea* (Xanthophycae, Fig 3 h), *Klebsormidium flaccidum* (Klebsormidiophyceae, Fig 3 h) and *Dicranum scopartium* (Bryophyta, Fig 3 h) peaked in July (245 days) or August (270 days). Relative abundances of *Microcoleus vaginatus* (Fig 3 h) and Oxalobacteriaceae (Proteobacteria, Fig 3 e) were elevated in surface soil relative to bulk soil across the entire period from January (70 days) through to August (270 days).

### Community assembly processes

There were differences in the community assembly processes which operated at the soil surface and in the bulk soil, across time and between microbial groups (Fig 4).

**Figure 4.**
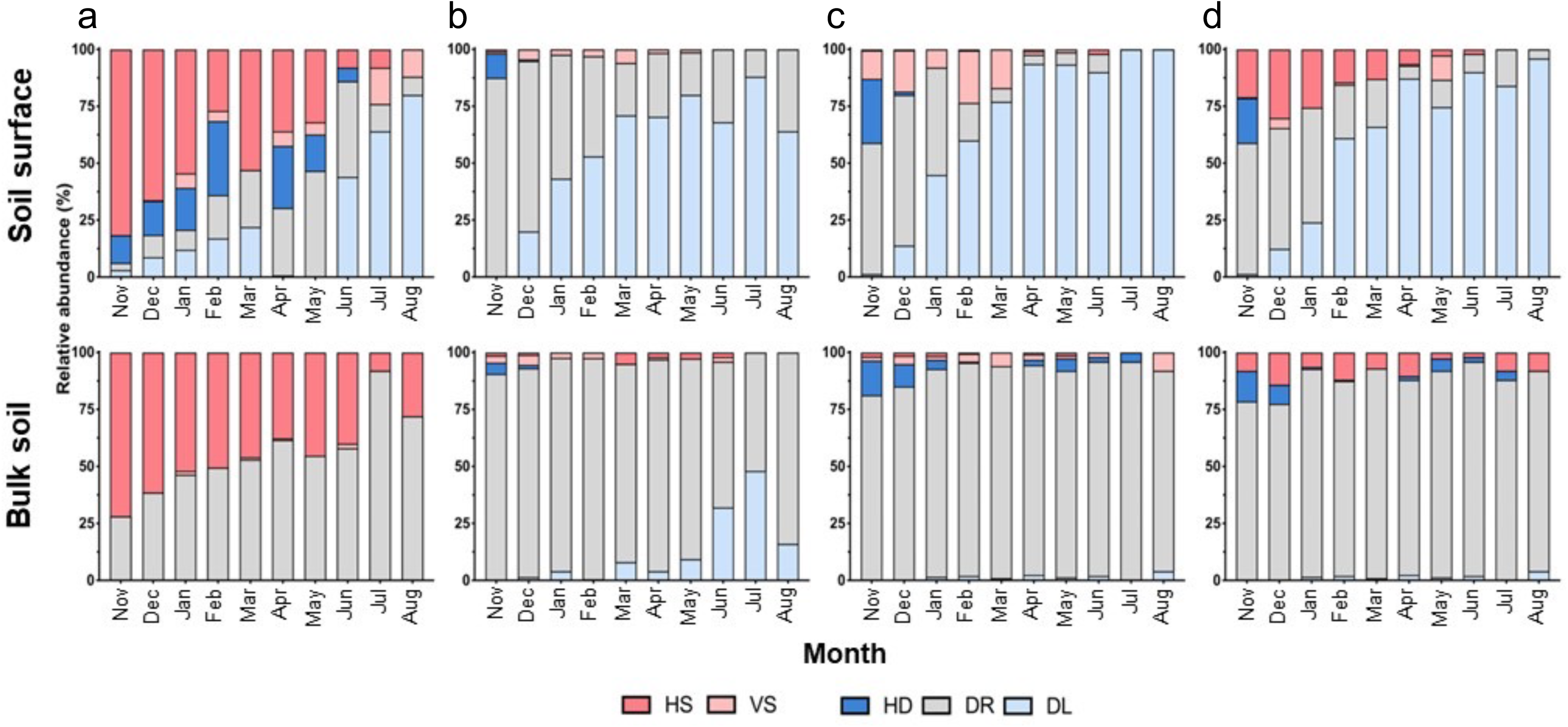
Community assembly processes in soil surface and bulk soil bacterial, fungal, protist and phototroph communities across time. a. Bacterial, b. Fungal, c. Protist, and d. Phototroph communities HS (homogenous selection) - β-NTI > 2, VS (variable selection) β-NTI < −2, HD (homogenising dispersal) β-NTI <|2| and B_RC_ > 0.95, DR (drift), and B_RC_ < |0.95| and DL and B_RC_ < −0.95 (dispersal limitation).

Bacterial communities in both the surface and bulk soil were initially dominated by homogenous selection, the importance of which declined over time, particularly in the surface soil (Fig 4 a). At the last sampling points in July (245 days) and August (270 days), variable selection was the most important deterministic selection process in surface soil but was of minor importance in bulk soil. Homogenous dispersal was not important in bulk soil but dominated stochastic processes in surface soil between November (0 days) and February (100 days), after which drift became increasingly important until June (220 days), when dispersal limitation became the dominant assembly process. In contrast, the importance of drift increased over time in the bulk soil as homogenous selection declined, with no evidence for dispersal limitation.

For fungi, drift was the most important assembly process initially, and its importance declined rapidly at the soil surface, as dispersal limitation became increasingly important over time (Fig 4 b). In contrast, drift remained the dominant assembly process in bulk soil through the entire time course. Homogenous dispersal was detected only at the initial November timepoint, with greater importance in surface relative to bulk soil. In protists, similar patterns to fungi were seen in the relative importance and dynamics of drift and dispersal limitation between the soil surface and bulk soil, and across time, although in the bulk soil community assembly was almost entirely attributed to drift (Fig 4 c). Variable selection was a contributor to community assembly between November and March (130 days) in the surface soil but had a minor contribution in bulk soil. Homogenous dispersal was the dominant stochastic process in the bulk and surface soil, with greater importance in the surface relative to the bulk soil at the initial November time point, although subsequently it contributed to assembly only in the bulk soil.

For phototrophs, the contribution of dispersal limitation to community assembly in surface soil increased over time, as drift declined, so that it dominated at the final August (270 days) time point (Fig 4 d). In bulk soil dispersal limitation made very little contribution to assembly. Homogenous selection and homogenous dispersal were more important contributors to assembly in surface soil than bulk soil at the initial November time point. After December (30 days), homogenizing dispersal was detected only in bulk soil, where it persisted in variable importance until July (245 days). While homogenizing selection declined in importance in surface soil over time, and disappeared in July (245 days), it remained a consistent but minor contributor to community assembly in bulk soil until August (270 days).

### Metagenomic comparison of surface and bulk soil microbial communities

Comparative metagenomic analysis indicated significantly higher (p < 0.01) relative abundance of carbon fixation genes in surface relative to bulk soil (Supplementary Fig 5 a). There was an associated significant (p < 0.01) enrichment in photosystem I and II (Fig 5 d, e), cytochrome b8f (Fig 5 c) and Rubisco (Fig 5 g). Across these processes, eukaryotes, including chlorophyta and streptophyta, and to a lesser extent cyanobacteria, were enriched in surface relative to bulk soil. Furthermore, the surface soil showed significantly (p < 0.01) greater abundance of the 3-hydroxypropionate bicycle (Fig 5 a) and the reductive TCA cycle CO_2_ fixation processes (Fig 5 f).

**Figure 5.**
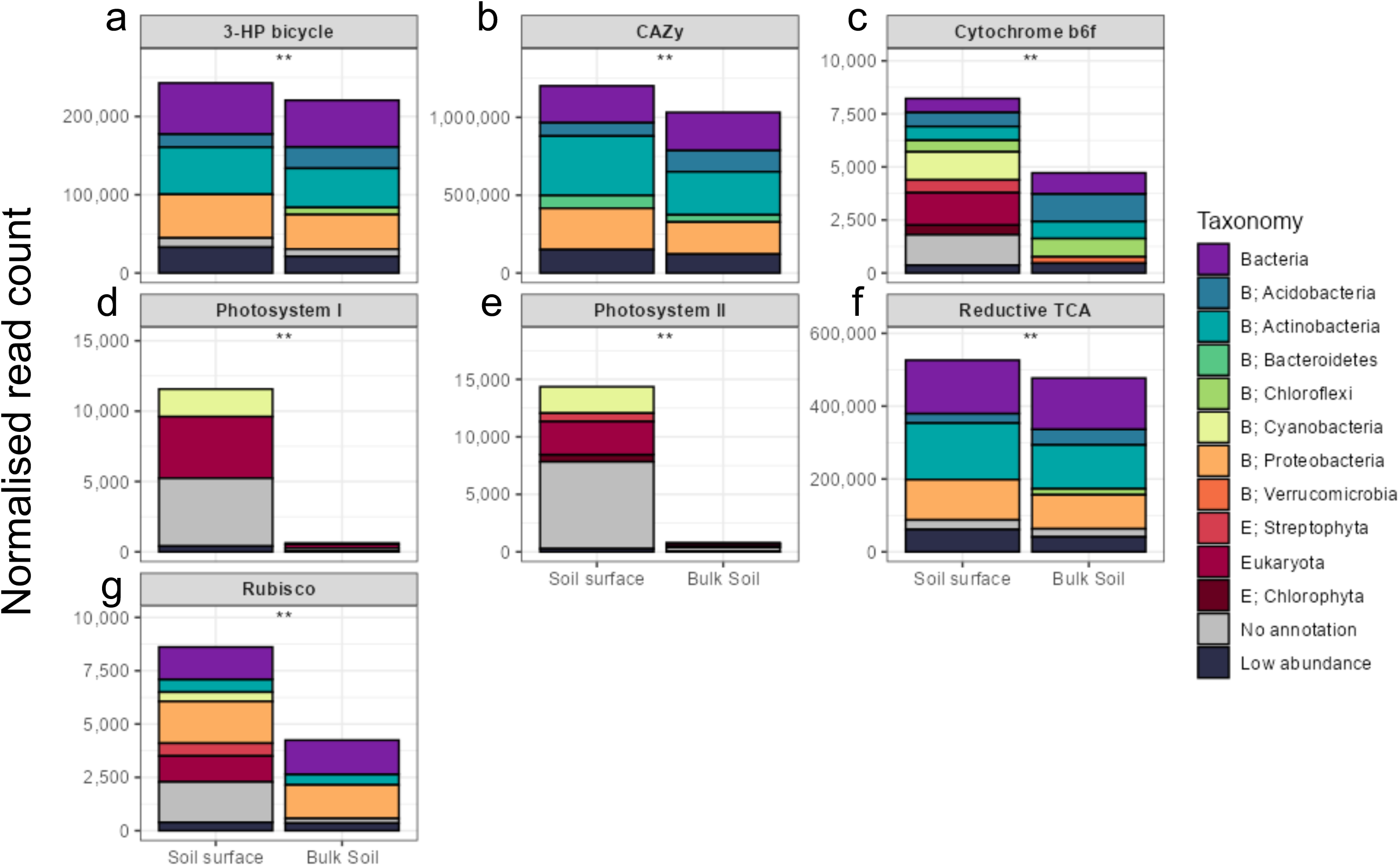
Normalised abundance taxonomic composition of of carbon cycle genes in the metagenome of soil surface and bulk soil communities. a. 3-hydroxypropionate bicycle, b. Carbohydrate active enzymes (CAZy), c. Cytochrome b6f, d. Photosystem I, e. Photosystem II, f. Reductive tricarboxylic acid (TCA) cycle, g.Ribulose-1,5-bisphosphate carboxylase/oxygenase (RUBISCO). Significance of difference in normalised read count between surface and bulk soil is indicated. **, significant p < 0.01.

There was also significantly (p < 0.01) elevated abundance of CAZy in surface relative to bulk soil (Fig 5 b) linked to significant (p < 0.01) enrichment of glycoside hydrolyase (Fig 6 a) and polysaccharide lyase (Fig 6 c), which are associated with degradation of plant materials such as pectin and cellulose. There was also significant (p < 0.05) enrichment of glycosyl transferase in surface soil (Fig 6 b), and this could reflect greater biosynthesis of disaccharides, oligosaccharides and polysaccharides associated with plant and microbial growth at the soil surface. For all of these genes, and particularly for polysaccharide lyase, increased abundance at the soil surface was associated with a marked increase in reads from Proteobacteria and Bacteroidetes.

**Figure 6.**
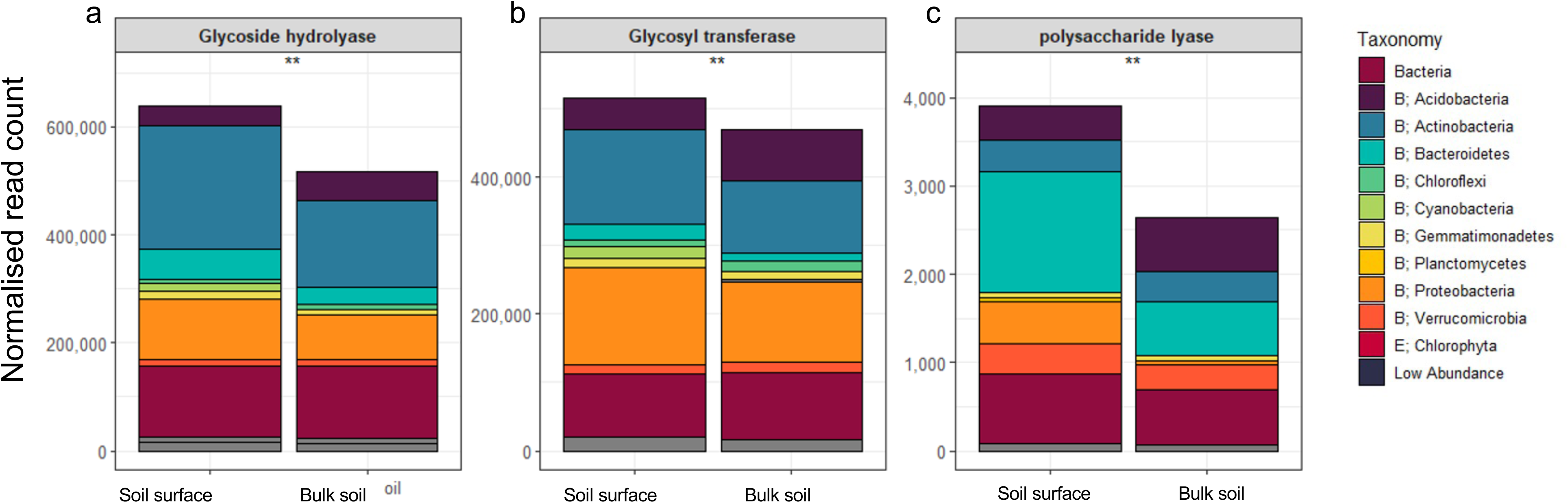
Normalised abundance and taxonomic composition of CAZy genes in the metagenome of surface and bulk soil. a. Glycoside hydrolyase, b. Glycosyl transferase, c. Polysaccharide lyase. Significance of difference in normalised read count between soil surface and bulk soil is indicated. **, significant p < 0.01.

Nitrogen cycle pathways were significantly (p < 0.01) enriched in the surface relative to bulk soil (Supplementary Fig 5 b). This included significantly (p < 0.01) increased relative abundance of denitrification (Fig 7 b), dissimilatory nitrate reduction (Fig 7 c), and nitrification pathways (Fig 7 d), largely attributed to increased abundance of Actinobacteria. Additionally, there was a significant (p < 0.01) increase in relative abundance of N_2_ fixation genes at the surface (Fig 7 e), relative to the bulk soil, and this was associated with cyanobacteria and proteobacteria.

**Figure 7.**
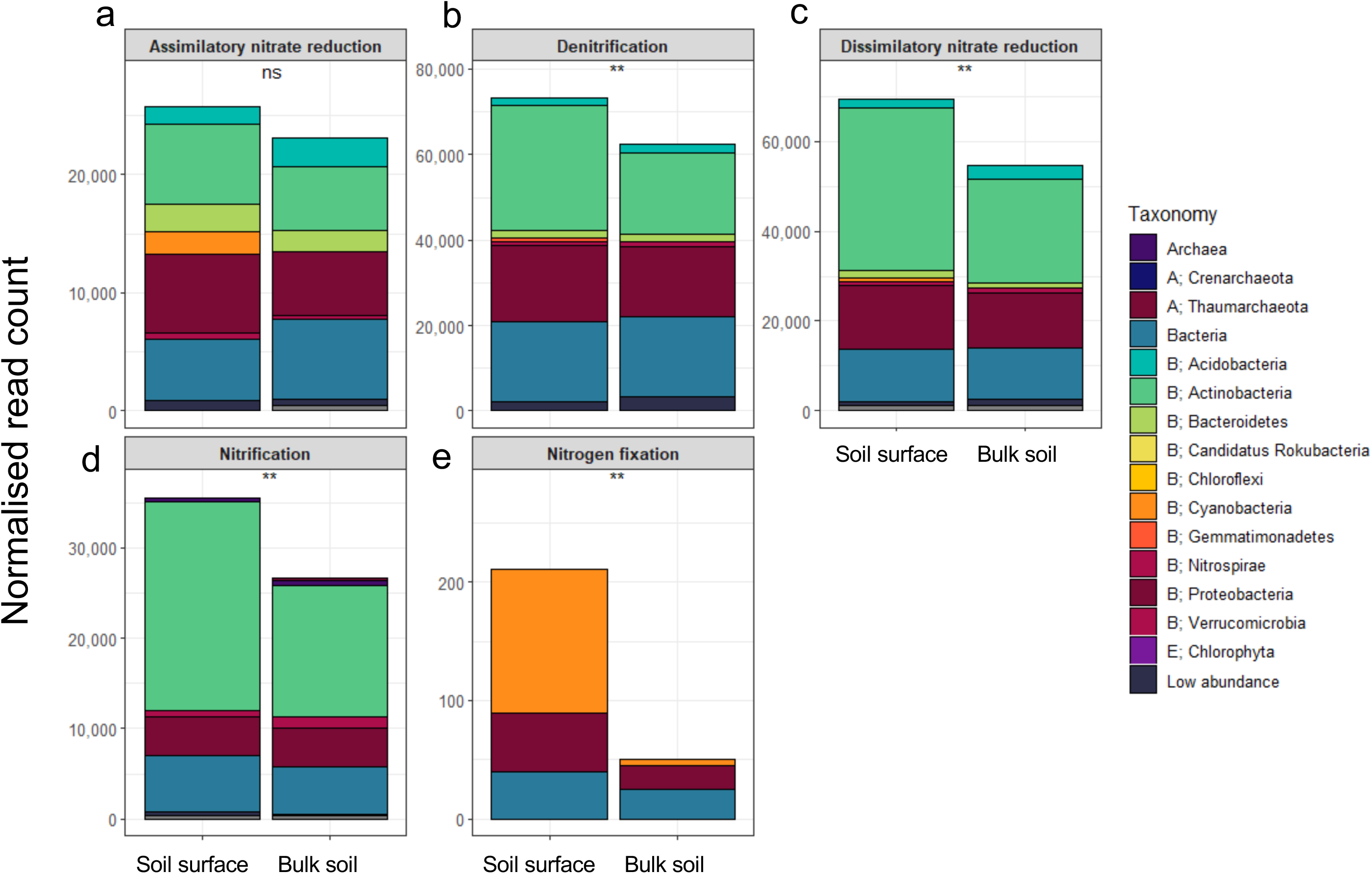
Normalised abundance and taxonomic composition of nitrogen cycling genes in the metagenome of soil surface and bulk soil. a. Assimilatory nitrate reduction, b. Denitrification, c. Dissimilatory nitrate reduction, d. Nitrification, e. Nitrogen fixation. Significance of differences in normalised read count between soil surface and bulk soil is indicated. ns, not significant; **, significant p < 0.01.

The soil surface also possessed significantly (p < 0.01) increased relative abundance of sulphur cycling genes relative to bulk soil (Supplementary Fig 5 c). Assimilatory and dissimilatory sulphate reduction showed significantly (p < 0.01) elevated relative abundance at the surface relative to the bulk soil, linked to increased abundance of genes from Actinobacteria, Bacteroidetes and Proteobacteria (Supplementary Fig 6 a, b). In contrast relative abundance of thiosulphate oxidation significantly (p < 0.01) reduced in the soil surface relative to the bulk soil and this reflected a reduction in genes from Acidobacteria at the soil surface (Supplementary Fig 6 c).

While there was no significant difference in relative abundance of P cycling pathways between the surface and bulk soil (Supplementary Fig 5 d), phosphatases (Supplementary Fig 7 a) and pathways associated with phosphate starvation (Fig 7 b), were enriched in the soil surface relative to bulk soil, and this reflected significantly (p < 0.01) greater abundance of genes from Proteobacteria, Actinobacteria and Bacteroidetes at the soil surface. There was no significant difference in abundance of phosphonatases, phytases or P transporters between surface and bulk soil.

## Discussion

We show rapid assembly of a taxonomically and functionally distinct soil surface microbiome in a tilled temperate agricultural soil, which involves succession of both phototrophic and heterotrophic communities over a 9-month period. Yellow-green algae were the initial phototrophic colonisers of the soil surface before proliferation of cyanobacteria and charophytes and finally bryophytes. Heterotrophic bacteria, fungal and protist taxa also showed distinct successional patterns as the surface community developed. Surface and bulk soil microbial communities were shaped by different assembly processes, and for surface communities these changed over time, with drift and selection dominating in early stages of development, before dispersal limitation became the dominant assembly process for all groups after 9 months. Metagenome analysis showed that after 9 months of development the soil surface was associated with greater abundance of genes associated with heterotrophic C, N, P and S biogeochemical cycling processes, relative to bulk soil, and this was linked to increased abundance of Actinobacteria, Bacteroidetes and Proteobacteria.

At the start of the sampling period, tillage ensured that there were no differences in community composition between surface and bulk soils. In the early months molecular evidence demonstrated development of a soil surface community associated with increased abundance of charophyte green algae (Klebsormidiophyceae), yellow-green algae (Xanthophyceae) and cyanobacteria (*Phormidium* and *Microcoleus*), which started prior to visible changes in soil surface morphology. These communities remained abundant through to the end of the monitoring period, although notably there were changes within the composition of the yellow green algae, with *Vaucheria* dominating in the summer period. *Klebsormidium, Phormidium* and *Microcoleus* are widely reported as components of BSC in arid soils (Veluci et al., 2006), indicating similarity in biological composition between BSC in arid and temperate soils. These taxa excrete mucilage and exopolysaccharide in arid soils and are important in binding and stabilising the soil surface (Garcia-Pichel, 2023), therefore they could perform a similar role in temperate soils.

Yellow-green algae do not appear to be detected in BSC within arid soils, although they have been isolated from temperate soils (Broady, 1979) and have been noted to form biological soil crusts in small, fast drying puddles within the Alps (Turk and Gartner, 2003). These phototrophs could therefore be distinct components of BSC in more hydric climate zones. Our data indicates that this group may be important members of the BSC right across the year, including the dry summer period. Yellow-green algae are filamentous, and can form mats, and these characteristics could provide similar roles in binding and stabilising the soil surface as cyanobacteria and charophytes. Notably, Bryophyta, including *Dicranum scoparium* increased abundance at the soil surface during the summer period. Mosses are considered slow colonisers of the soil surface which have been recorded at the soil surface of non-agricultural arid ecosystems after 5-10 years, or approximately 2 years in non-agricultural temperate ecosystems (Maestre et al., 2011; Dojani et al., 2011; Fischer et al., 2010), therefore moss development appears to have been very rapid.

Evidence also suggested temporal dynamics across non-phototrophic communities. For some heterotrophs this could reflect adaptation to a soil surface environment. The ascomycete fungus *Exophiala,* which showed elevated relative abundance at the soil surface have melanised hyphae (Revanker and Sutton, 2010). This may be an adaptation to a light exposed habitat, since melanin provides protection against the damaging effects of UV light on DNA (Singaravelan et al., 2008). Changes in many heterotrophs may relate to biological interactions, potentially related to altered abundance of phototrophs or other heterotrophs. The Amoebozoa, and the likely heterotrophic flagellates *Allantion* (Ekelund and Patterson, 1997) and *Proleptomonas* (Vickerman et al., 2002), showed increased relative abundance at the soil surface, and these shifts could reflect changes in abundance of bacteria on which these taxa likely predate. This could include cyanobacteria, but also heterotrophic bacteria including Actinobacteria, Proteobacteria and Bacteroidetes in which there were marked temporal shifts in relative abundance. In addition, Meliniomyces may be root or rhizoid associated fungi (Hambleton and Sigler, 2005), and their elevated abundance in the soil surface during the summer could be linked to increased growth of plants, most likely bryophytes and soil surface inhabiting Viridiplantae.

Other changes in relative abundance of biota at the soil surface could reflect biotic signatures deposited at the soil surface from above ground components of the ecosystem, rather than growth on the soil surface. In particular, changes to the fungal community were associated with increased abundance of plant pathogens over the spring and summer period, including the generalist plant pathogens *Plenodomus biglobosus* (de Gruyter et al., 2013) and *Cladosporium exasperatum* (Bensch et al., 2010) and *Zymoseptoria* sp. a widespread pathogen of the wheat overstorey crop (Torriani et al., 2015). Interestingly, Eimeriidae are a group of protist parasites which largely infect the intestines of vertebrates (Ghimire, 2010), and increased relative abundance of this group in the summer could be a signature of infected animals at the field site, including mammals such as rabbits and bats, and birds.

Community assembly in bulk soil was dominated by drift in all taxa, indicating low turnover rates and weak selection pressures relative to surface soil. Notably for bacteria, but not the other taxa, homogenous selection was an important process in both bulk and surface soils. This occurs in environments within which there is a consistent environmental selection pressure and similar community composition (Zhou and Ning, 2017). In the very early stages of surface and bulk soil community development, homogenising dispersal was detected in all taxa, indicating that a high dispersal rate generated similarity in community composition (Zhou and Ning, 2017). However, the dominant process operating at the soil surface across all groups of biota then switched increasingly to dispersal limitation which was largely absent in bulk soil. Dispersal limitation occurs when there are weak selection pressures and physical barriers constrain organisms from different populations from mixing (Zhou and Ning, 2017) and has been identified as a key determinant of assembly in dryland BSC (Li and Hu, 2021). This was associated with extensive establishment of biota at the soil surface, which could have provided a physical and biological barrier to the establishment of new populations, ensuring dominance of stochastic assembly processes.

Metagenome analysis demonstrated that the soil surface was enriched in a variety of functions related to photosynthesis in eukaryotic algae and cyanobacteria including Rubisco, photosystems I and II and cytochrome b6f. However, there was also enrichment of non-photosynthetic reductive TCA and 3HP-bicycle C fixation processes in the soil surface, similar to the situation in dryland BSC in which non-photosynthetic C fixation processes vary across successional stages (Wang et al., 2022).

The soil surface showed four-fold higher relative abundance of N_2_ fixation genes than the bulk soil, attributed to both cyanobacteria and proteobacteria, although abundance within the metagenome was low. N_2_ fixation is considered a key characteristic of phototrophic communities inhabiting the surfaces of rocks, plants and soil, which may be responsible for half of the N_2_ fixation in terrestrial systems (Elbert et al., 2012). While cyanobacterial N_2_ fixation is well known in arid soil BSC (Belnap, 2002), there is limited understanding of its significance in temperate zones. Witty et al (1979) measured up to 28 kg N ha^-1^ fixed in areas of a UK arable field in which the soil surface was visibly colonised by cyanobacteria. Subsequent experiments in which cyanobacteria were inoculated onto the soil surface demonstrated lower rates of N_2_ fixation (5 kg N ha^-1^), with rates markedly affected by soil moisture regime (Witty, 1979). Our data therefore supports the potential for N_2_ fixation by soil surface biota in temperate agricultural systems and suggests that management of soil surface biota could play an important role the development of sustainable systems in which energy intensive and environmentally damaging fertiliser N inputs are reduced.

With the exception of C and N fixation, changes in heterotrophic, rather than phototrophic communities were largely responsible for changes in abundance of biogeochemical cycling genes between surface and bulk soil communities. Actinobacterial genes were particularly important in driving these differences for the CAZy enzymes glycoside hydrolase and glycosyl transferase, assimilatory and dissimilatory sulphate reduction, denitrification, dissimilatory nitrate reduction, nitrification, and also phosphatases. Bacteroidetes and Proteobacteria also contributed to greater abundance of genes in the soil surface for assimilatory and dissimilatory SO_4_ reduction, phosphatases, PO_4_ starvation, and polysaccharide lyase. While the importance of the soil surface as significant zone of biogeochemical cycling in arid regions is recognised, largely because of its role in fixation of C and N (Belnap, 2002, Elbert, 2012), our results indicate that the same is true in temperate agricultural areas, but moreover that the soil surface communities are zones of broad importance for C, N, P and S biogeochemical cycling by both heterotroph and phototroph communities.

The increased abundance of polysaccharide lyase genes, which cleave a range of substrates including pectin and glycosaminoglycans, and glycoside hydrolase genes, which break down cellulose and hemicellulose, at the soil surface relative to bulk soil could reflect increased heterotroph activity associated with decomposition of C returned to the soil surface by phototroph communities as exuded polysaccharides and biomass. However, inputs of C by phototrophs could also prime decomposition of native soil organic materials by heterotrophs (Henneron et al., 2020). This could also explain the increased abundance of genes associated with N, P and S cycling pathways in the soil surface relative to the bulk soil. In this way the interaction between heterotrophs and phototrophs at the soil surface could be analogous to interaction between plants and heterotrophic rhizosphere biota (Davies et al., 2013; Couradeau et al., 2019).

While the potential for management of soil surface biota for the restoration of degraded drylands is recognised (Coban et al., 2022), our data indicates that similar phototrophic and heterotrophic communities inhabit the surface of temperate agricultural soils, and that these biota could play vital roles in protecting the soil surface against erosion, which is a key challenge in agricultural systems, and will become more so as climate change exposes the soil surface to extreme weather (Li and Fang, 2016). Furthermore, C and N fixation by soil surface phototroph communities, and the resulting stimulation of heterotrophic biogeochemical cycling processes points to the soil surface as a dynamic compartment within temperate agricultural soil systems which is currently overlooked. Effective management of soil surface biota could therefore play a critical role in enhancing soil sustainability through soil protection, promoting C drawdown from the atmosphere to contribute to climate mitigation, and reducing the need for mineral fertiliser inputs.

## Acknowledgements

We thank the Natural Environment Research Council and the Biotechnology and Biological Sciences Research Council for funding

