## Supplementary Figures for "Rapid Assembly and Functional Differentiation of the Soil Surface Microbiome in Temperate Agricultural soil"

### Slide 1
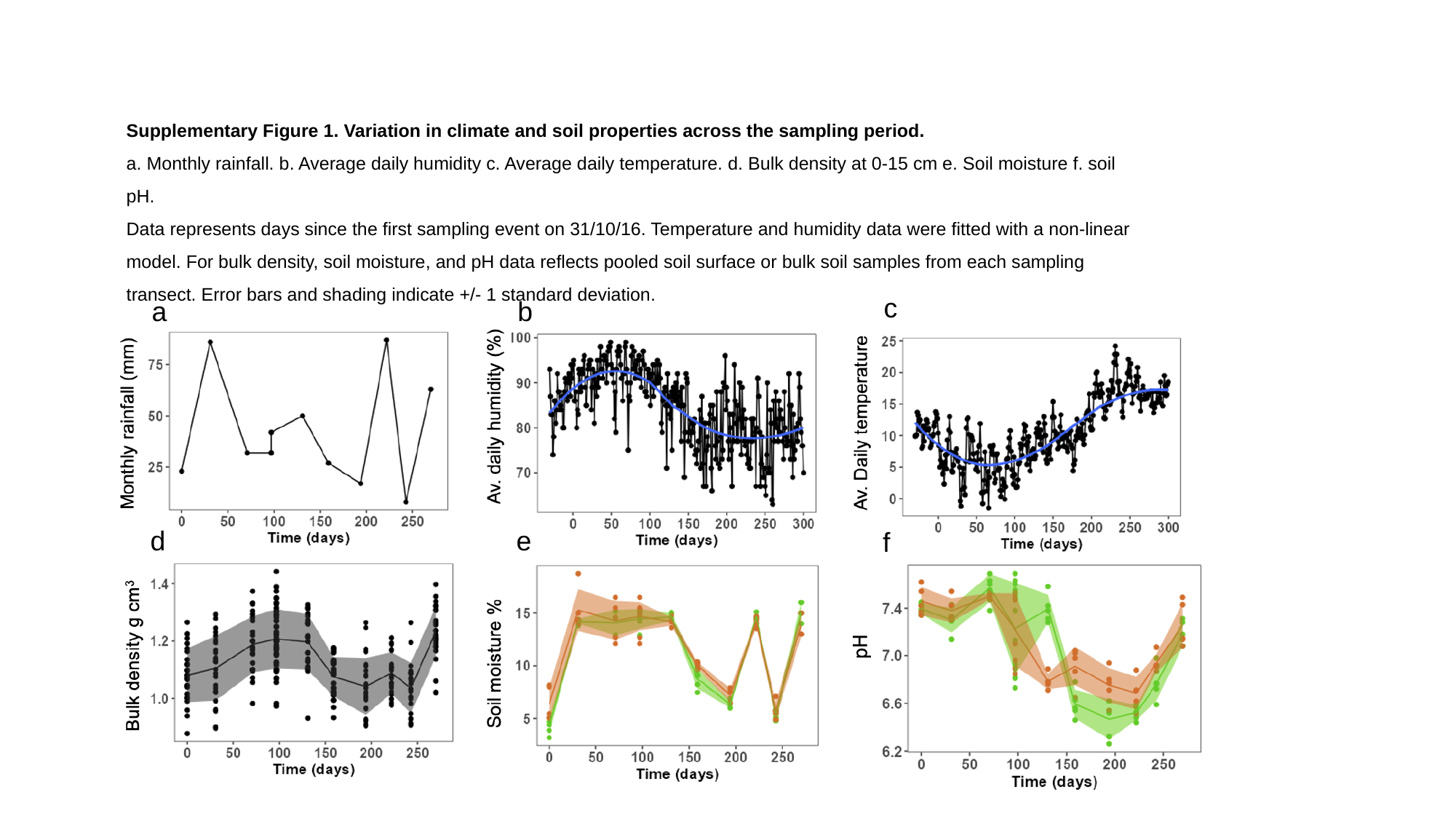

Supplementary Figure 1. Variation in climate and soil properties across the sampling period.
a. Monthly rainfall. b. Average daily humidity c. Average daily temperature. d. Bulk density at 0-15 cm e. Soil moisture f. soil pH.
Data represents days since the first sampling event on 31/10/16. Temperature and humidity data were fitted with a non-linear model. For bulk density, soil moisture, and pH data reflects pooled soil surface or bulk soil samples from each sampling transect. Error bars and shading indicate +/- 1 standard deviation.
c
b
a
e
d
f

### Slide 2
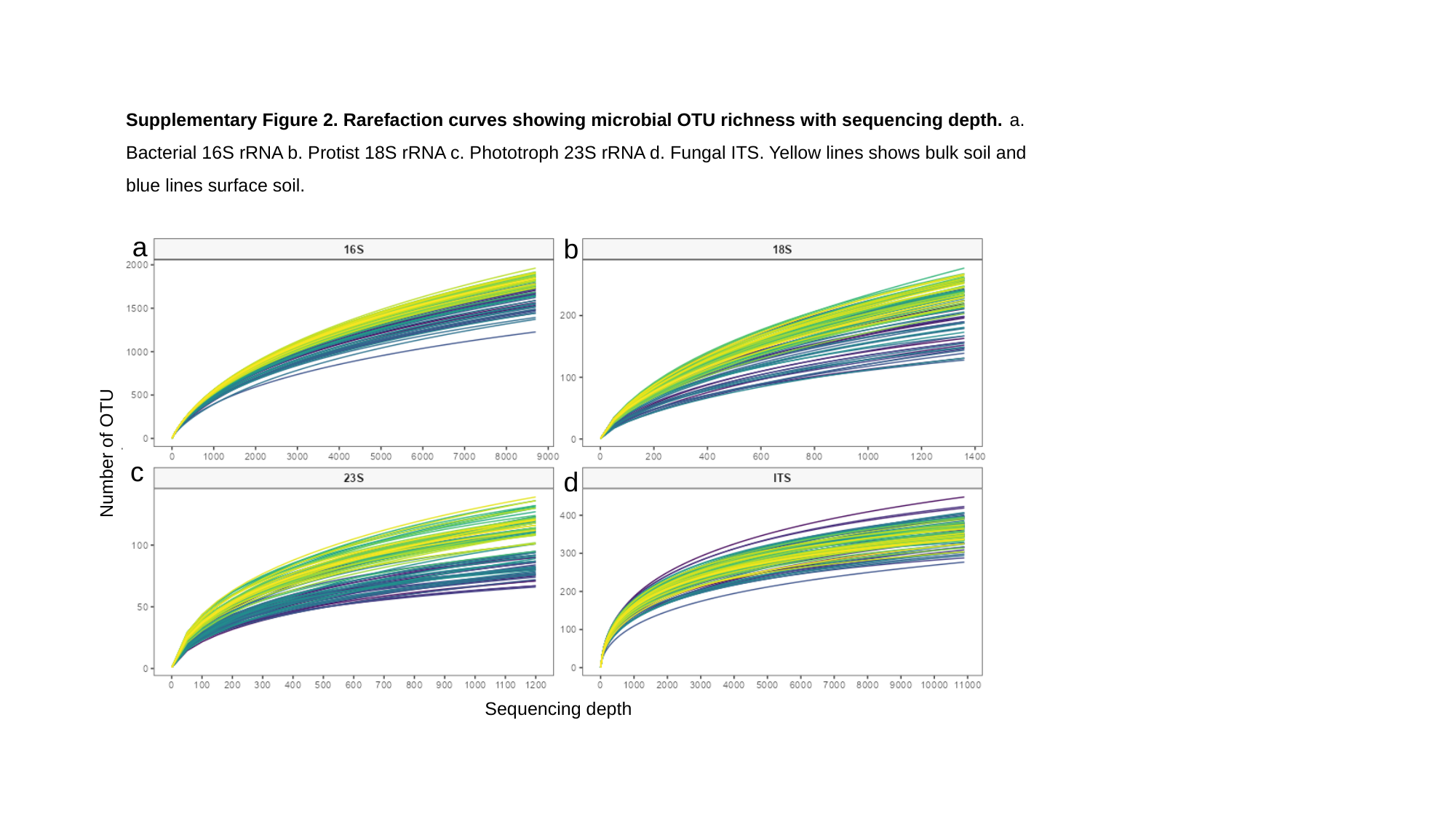

Supplementary Figure 2. Rarefaction curves showing microbial OTU richness with sequencing depth. a. Bacterial 16S rRNA b. Protist 18S rRNA c. Phototroph 23S rRNA d. Fungal ITS. Yellow lines shows bulk soil and blue lines surface soil.
a
b
Number of OTU
c
d
Sequencing depth

### Slide 3
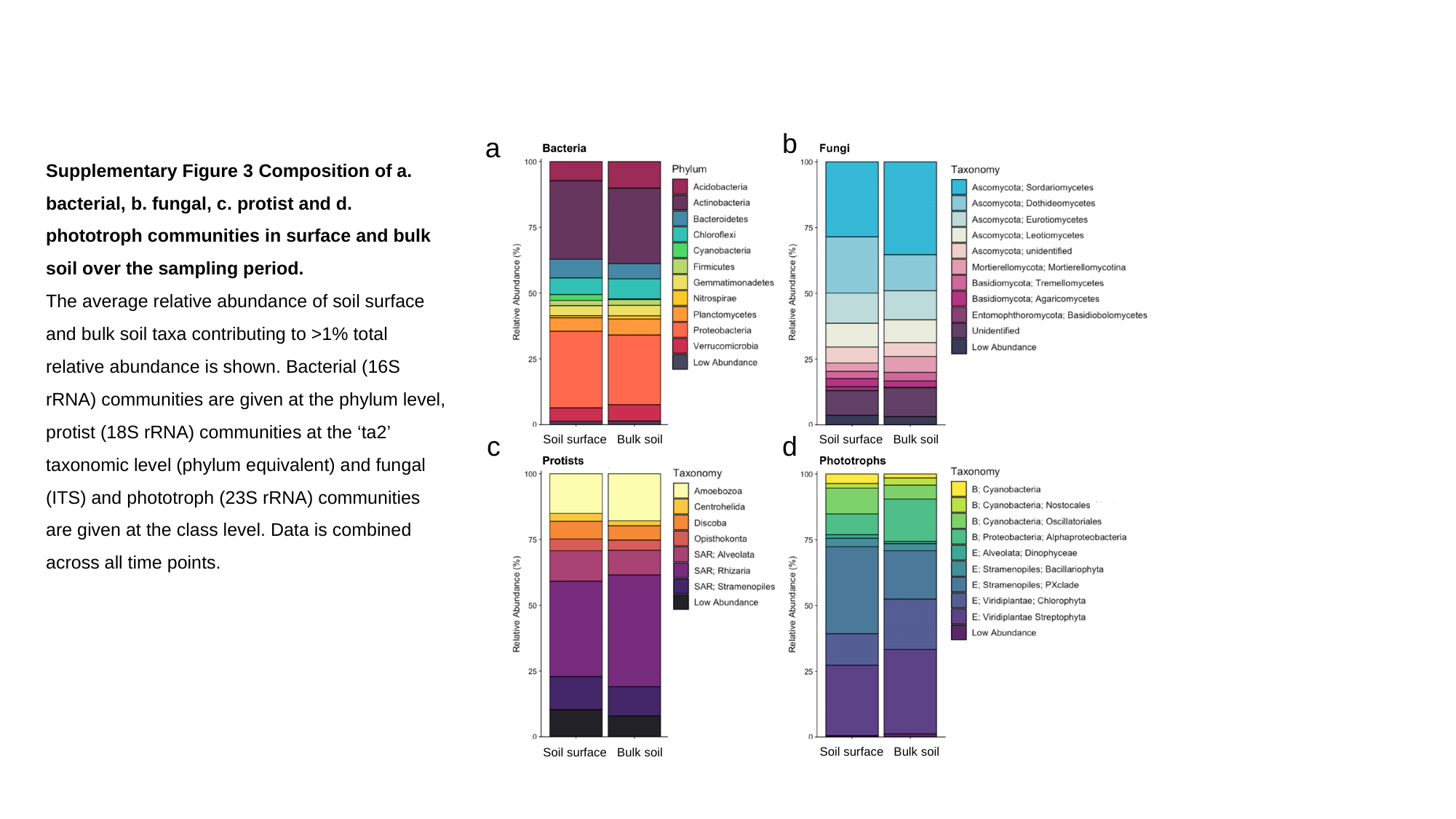

b
a
Supplementary Figure 3 Composition of a. bacterial, b. fungal, c. protist and d. phototroph communities in surface and bulk soil over the sampling period.
The average relative abundance of soil surface and bulk soil taxa contributing to >1% total relative abundance is shown. Bacterial (16S rRNA) communities are given at the phylum level, protist (18S rRNA) communities at the ‘ta2’ taxonomic level (phylum equivalent) and fungal (ITS) and phototroph (23S rRNA) communities are given at the class level. Data is combined across all time points.
c
d
Soil surface Bulk soil
Soil surface Bulk soil
Soil surface Bulk soil
Soil surface Bulk soil

### Slide 4
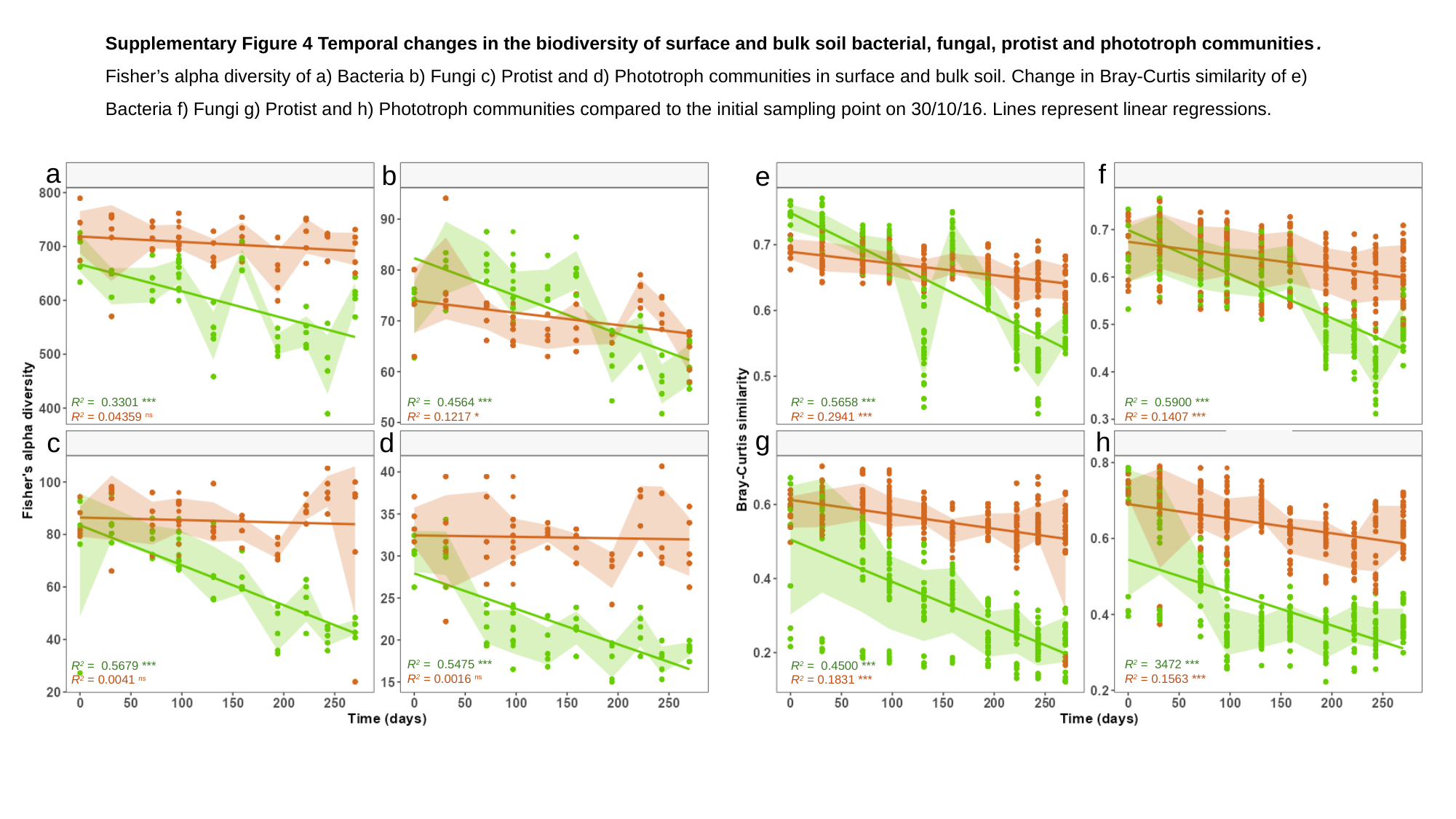

Supplementary Figure 4 Temporal changes in the biodiversity of surface and bulk soil bacterial, fungal, protist and phototroph communities.
Fisher’s alpha diversity of a) Bacteria b) Fungi c) Protist and d) Phototroph communities in surface and bulk soil. Change in Bray-Curtis similarity of e) Bacteria f) Fungi g) Protist and h) Phototroph communities compared to the initial sampling point on 30/10/16. Lines represent linear regressions.
a
f
b
e
R^2^ = 0.3301 ***
R^2^ = 0.04359 ^ns^
R^2^ = 0.4564 ***
R^2^ = 0.1217 *
R^2^ = 0.5658 ***
R^2^ = 0.2941 ***
R^2^ = 0.5900 ***
R^2^ = 0.1407 ***
g
h
c
d
R^2^ = 0.5475 ***
R^2^ = 0.0016 ^ns^
R^2^ = 3472 ***
R^2^ = 0.1563 ***
R^2^ = 0.5679 ***
R^2^ = 0.0041 ^ns^
R^2^ = 0.4500 ***
R^2^ = 0.1831 ***

### Slide 5
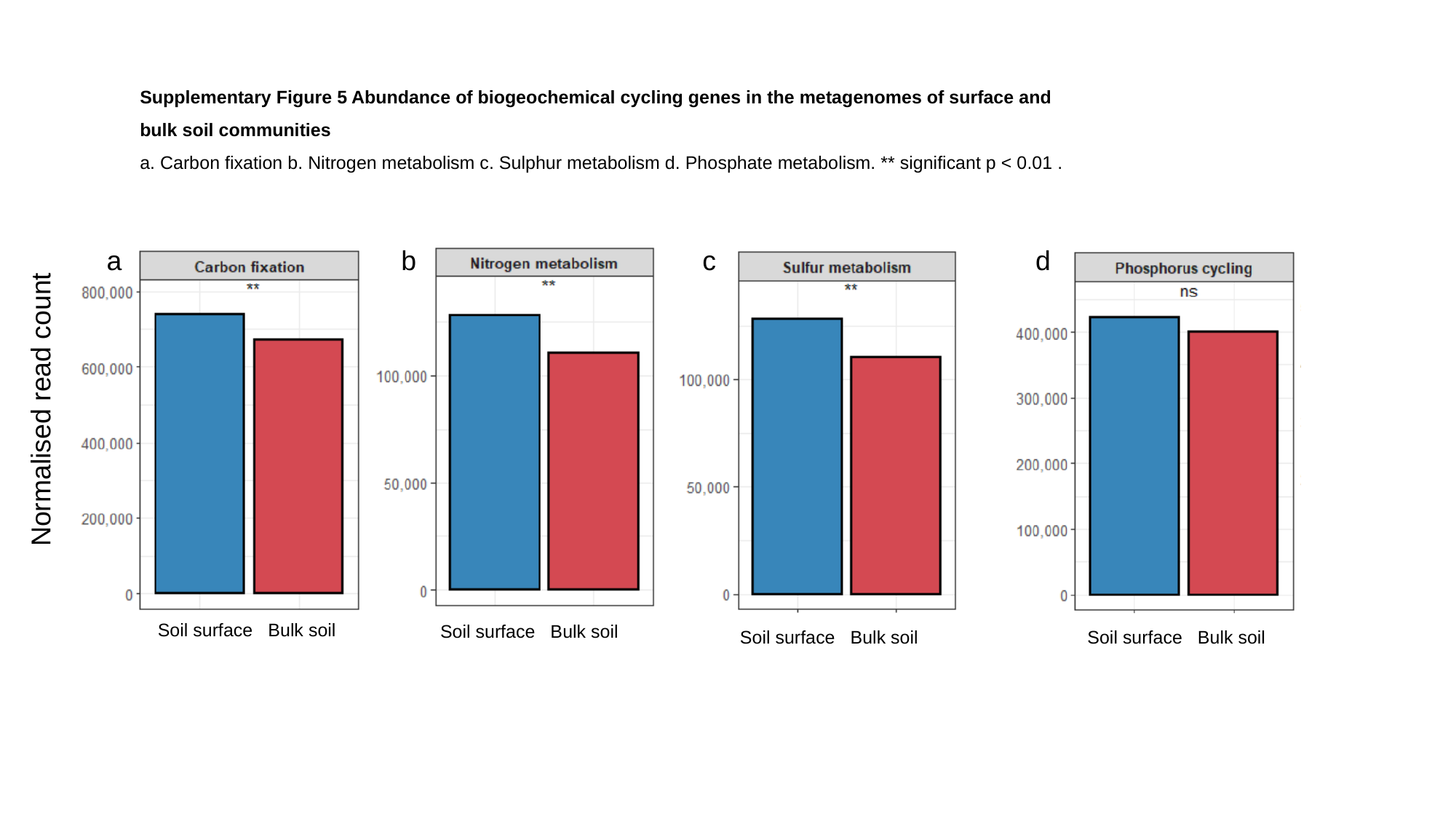

Supplementary Figure 5 Abundance of biogeochemical cycling genes in the metagenomes of surface and bulk soil communities
a. Carbon fixation b. Nitrogen metabolism c. Sulphur metabolism d. Phosphate metabolism. ** significant p < 0.01 .
a
b
c
d
Normalised read count
Soil surface Bulk soil
Soil surface Bulk soil
Soil surface Bulk soil
Soil surface Bulk soil

### Slide 6
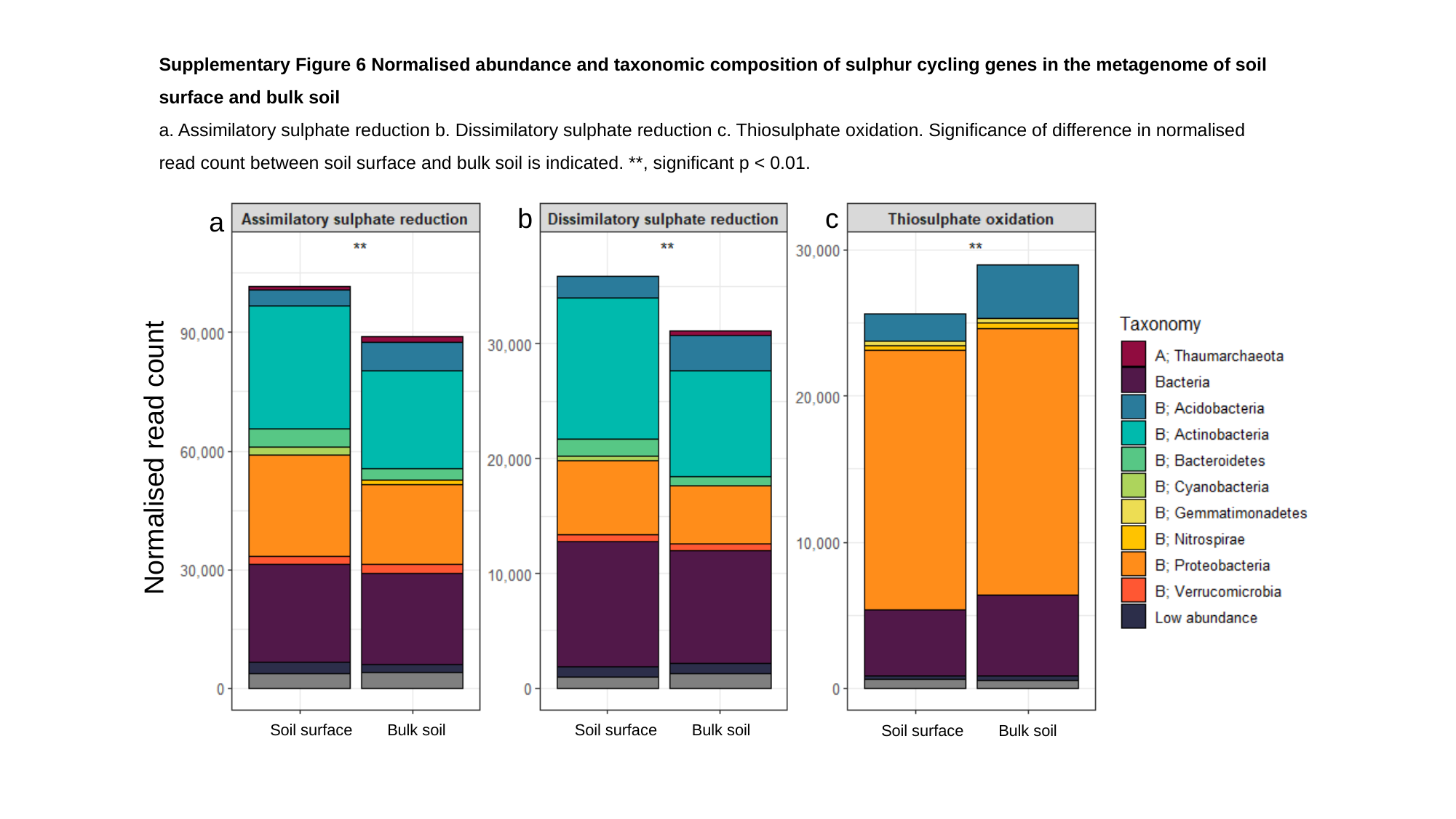

Supplementary Figure 6 Normalised abundance and taxonomic composition of sulphur cycling genes in the metagenome of soil surface and bulk soil
a. Assimilatory sulphate reduction b. Dissimilatory sulphate reduction c. Thiosulphate oxidation. Significance of difference in normalised read count between soil surface and bulk soil is indicated. **, significant p < 0.01.
b
c
a
Normalised read count
Soil surface Bulk soil
Soil surface Bulk soil
Soil surface Bulk soil

### Slide 7
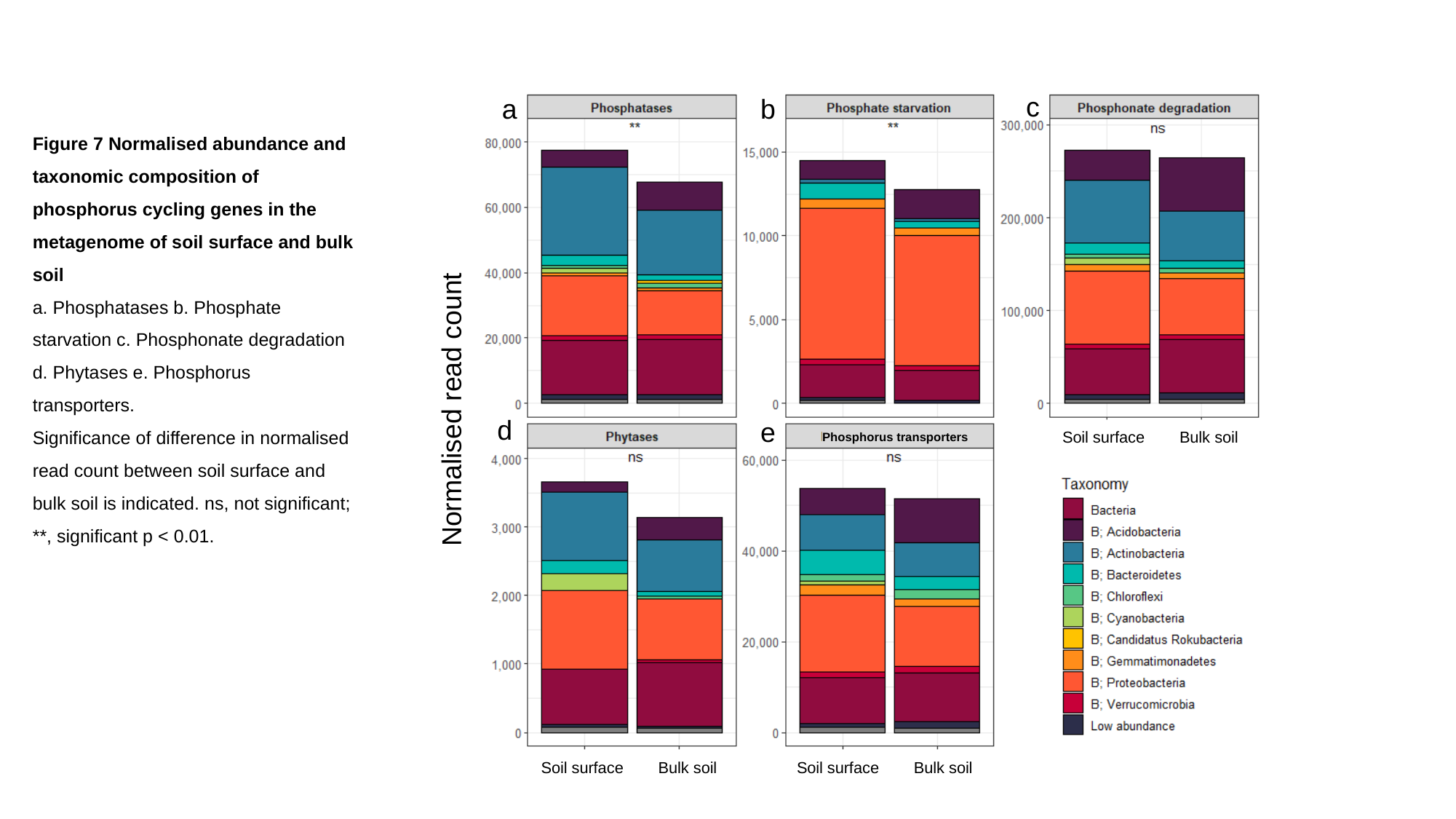

c
a
b
Figure 7 Normalised abundance and taxonomic composition of phosphorus cycling genes in the metagenome of soil surface and bulk soil
a. Phosphatases b. Phosphate starvation c. Phosphonate degradation d. Phytases e. Phosphorus transporters.
Significance of difference in normalised read count between soil surface and bulk soil is indicated. ns, not significant; **, significant p < 0.01.
Normalised read count
d
e
Soil surface Bulk soil
Phosphorus transporters
Soil surface Bulk soil
Soil surface Bulk soil
