## Supplementary Table 1 for "Rapid Assembly and Functional Differentiation of the Soil Surface Microbiome in Temperate Agricultural soil"

Table S1 Similarity percentage (SIMPER) analysis of OTU contributing to dissimilarity of soil surface communities over time. Shown are the 5 OTUs with the largest contributions to community dissimilarity across sampling times in the surface soil.

| Taxon | OTU | | Av. dissimilarity | % contribution |
| --- | --- | --- | --- | --- |
| **Bacteria** | |  |  |  |
| Oxalobacteraceae | | OTUB2 | 0.664 | 1.65 |
| Pseudomonas | | OTUB31 | 0.654 | 1.62 |
| Phormidium | | OTUB15 | 0.569 | 1.41 |
| Micrococcaceae | | OTUB1 | 0.414 | 1.03 |
| Flavobacterium | | OTUB32 | 0.340 | 0.84 |
| **Fungi** | |  |  |  |
| Plenodomus biglobosus | | OTUF16 | 2.12 | 4.79 |
| Cladosporium exasperatum | | OTUF2 | 1.70 | 3.86 |
| Exophiala equina | | OTUF1 | 1.32 | 3.00 |
| Zymoseptoria brevis | | OTUF27 | 1.14 | 2.58 |
| Meliniomyces | | OTUF25 | 1.08 | 2.44 |
| **Protists** | |  |  |  |
| Allantion | | OTUP1569 | 8.18 | 13.02 |
| Eimeridae | | OTUP1561 | 4.31 | 6.86 |
| Retaria | | OTUP2738 | 1.51 | 2.40 |
| Proleptomonas faecicola | | OTUP1188 | 1.28 | 2.03 |
| Chromulinales | | OTUP1562 | 1.11 | 1.77 |
| **Phototrophs** | |  |  |  |
| Vaucheria littorea | | OTUPh2 | 10.18 | 18.9 |
| Dicranium scoparium | | OTUPh5 | 4.02 | 7.46 |
| Klebsormidium flaccidum | | OTUPh1 | 3.97 | 7.37 |
| Microcoleus vaginatus | | OTUPh1718 | 2.39 | 4.44 |
| Acetobacteriaceae | | OTUPh3 | 1.85 | 3.44 |
